# Optimization of Gadolinium-Based Contrast Agent Protocols for Reliable Ex Vivo Diffusion-Weighted Imaging in the Avian Brain

**DOI:** 10.64898/2026.06.19.733394

**Authors:** Marie Ziegler, Paula Gerliz, Xavier Helluy, Onur Güntürkün, Mehdi Behroozi

**Author notes:** Corresponding Author: Marie Ziegler.

## Abstract

Ex vivo diffusion weighted imaging (DWI) enables high-resolution characterization of brain connectivity and is increasingly applied in comparative and evolutionary neuroscience. However, variability in tissue preparation and contrast agent exposure can substantially affect relaxation properties and compromise reproducibility, particularly in non-mammalian species. Here, we systematically assess the impact of different gadolinium-based contrast agent exposure protocols on relaxation stability and DWI compatibility in fixed pigeon brains.

Brains were perfusion-fixed with 2% paraformaldehyde and assigned to four preparation protocols: (i) contrast agent exposure during perfusion, post-fixation, and rehydration; (ii) post-fixation and rehydration only; (iii) rehydration only; (iv) no contrast agent. Quantitative T1, T2, T2*, and DWI data were acquired at five time points over 70 days using a 7T MRI system.

Protocols involving contrast agent during perfusion or post-fixation produced comparable relaxation trajectories, with T1, T2, and T2* stabilizing by Day 13. On day 13 the T1 values of tissue that was exposed to contrast agent, regardless of the application protocol were between 230.86 ms and 266.89 ms, while the T1 values of the control group were over 1100 ms at this point in time. T2 values of the experimental groups were between 39.97 ms and 56.17 ms while T2 values of the control group were between 58.68 ms and 77.82 ms. T2* values of the experimental groups were between 27.27 ms and 43.33 ms while T2* values of the control group were between 46.16 ms and 65.93 ms. Importantly, contrast agent exposure during rehydration alone resulted in equivalent stabilization after two weeks, reflecting gradual contrast agent diffusion into the tissue. In contrast, control samples without contrast agent exhibited significantly elevated T2 and T2* at later time points.

These results demonstrate that post-fixation contrast agent exposure during rehydration is sufficient to achieve stable relaxation parameters and DWI compatibility, assessed via fractional anisotropy (FA) and mean diffusivity (MD) in ex vivo avian brain tissue. This minimal preparation protocol enhances reproducibility, reduces handling complexity, and supports standardized cross-species neuroimaging of brain connectivity.

## 1 Introduction

Diffusion weighted imaging (DWI) is a magnetic resonance imaging (MRI) technique that allows the characterization of tissue microstructure by quantifying the directionality and magnitude of water diffusion. In neuroscience, DWI is used to map white matter pathways, reconstructing structural brain networks, and linking microstructural organization to development, learning and disease (Assaf et al., 2019; Basser, 1995; Blumenfeld-Katzir et al., 2011; Chien et al., 1992; Moseley et al., 1990; Qiu et al., 2015). While in vivo diffusion MRI is constrained by motion, scan time, and safety considerations, ex vivo DWI allows substantially longer acquisition and higher spatial resolution, providing unique opportunities for detailed anatomical investigation and cross-modal validation with histology (Behroozi et al., 2024; D’Arceuil et al., 2007; Dyrby et al., 2011; Pfefferbaum et al., 2004; Roebroeck et al., 2019).

However, despite the advantages, ex vivo diffusion imaging presents substantial technical challenges. Tissue fixation shortens T1 and T2 relaxation times and reduces diffusivity, with effects depending on fixative concentration, post-fixation handling, and rehydration times, ultimately decreasing the signal-to-noise ratio (SNR) and necessitating careful optimization of tissue preparation and acquisition parameters (Barrett et al., 2023; Blamire et al., 1999; D’Arceuil et al., 2007; Dawe et al., 2009; Pfefferbaum et al., 2004; Raman et al., 2017; Shepherd et al., 2009; Sun et al., 2005; Thelwall et al., 2006; Tovi & Ericsson, 1992). Raman et al. (2017) showed in their study, that formalin fixation leads to a substantial reduction in T1 relaxation times in ex vivo human brain tissue, in both white and grey matter during the first weeks of fixation. This initial rapid shortening of T1 was then followed by a further yet slower, decline over time. Dawe et al. (2009) also showed in ex vivo human brain tissue, that T2 relaxation times decrease as a result of formaldehyde fixation with the most pronounced decreases occurring in tissue regions exposed to the fixative early in the fixation process.

To mitigate these limitations, gadolinium-based contrast agents have been widely adopted in ex vivo MRI to reduce longitudinal (T1) relaxation times and to increase SNR efficiency, thereby enabling shorter repetition times (TR) and higher spatial resolution (Barrett et al., 2023; Benveniste & Blackband, 2002; D’Arceuil et al., 2007; Huang et al., 2009; Ullmann et al., 2010; Weinmann et al., 1984). However, inappropriate contrast agent concentrations or application protocol can excessively shorten transverse relaxation times (T2, T2*), degrade image quality, and bias diffusion metrics (Barrett et al., 2023; D’Arceuil et al., 2007; Huang et al., 2009; Ullmann et al., 2010; Weinmann et al., 1984). A study by D’Arceuil et al. (2007) investigating the use of gadolinium-based contrast agents in fixed brain tissue showed that gadolinium reduces T1 relaxation times and thereby improves scan efficiency. They also showed that the contrast agent altered tissue relaxation properties and influenced quantitative diffusion measurements, thereby underlining the importance of optimizing contrast agent concentrations and fixation protocols for ex vivo diffusion MRI (dMRI).

Previous work has demonstrated that systematic optimization of tissue preparation parameters, including fixative concentration, rehydration time, and contrast agent concentration, can substantially improve data quality in ex vivo diffusion MRI (Barrett et al., 2023; D’Arceuil et al., 2007; Dyrby et al., 2011; Roebroeck et al., 2019; Ullmann et al., 2010). In rodents and primates, reducing fixative concentration, allowing extended tissue equilibration, and carefully titrating gadolinium exposure have been shown to increase SNR efficiency while largely preserving diffusion tensor metrics such as fractional anisotropy (FA) and mean diffusivity (MD) (Barrett et al., 2023; D’Arceuil et al., 2007; Shepherd et al., 2009). While these studies provide an important conceptual framework for tissue optimization, they are mainly focused on mammalian brains. Therefore, the transferability to vertebrate systems with different brain organization and tissue composition remains largely unexplored.

Avian brains present a particularly compelling yet underexplored target for high-resolution ex vivo diffusion imaging. Birds possess highly specialized neural circuits supporting spatial navigation (Mouritsen et al., 2016; Rook et al., 2023), visual processing (Heyers et al., 2008; Pusch et al., 2023; Rook, Tuff, Isparta, et al., 2021), and cognitive flexibility (Emery & Clayton, 2004; Nieder, 2017; Rook, Tuff, Packheiser, et al., 2021; Steinemer et al., 2024). Furthermore, they show remarkable differences from mammals in pallial organization, nuclear architecture, and white matter arrangement (Güntürkün & Bugnyar, 2016; Jarvis et al., 2005; Jarvis et al., 2013; Stacho et al., 2020). These features make birds an interesting model for comparative and evolutionary neuroscience, however they also pose unique challenges for dMRI. Differences in tissue relaxation behavior, and diffusion stability, make mammalian optimization protocols suboptimal or inappropriate for avian tissue. To date, the influence of contrast agent application timing on relaxation properties and diffusion metrics in the avian brain remains unexplored.

In this study, we systematically evaluate gadolinium-based contrast agent application strategies for ex vivo DWI of the pigeon brain, with the goal to identify a protocol that balances signal enhancement while preserving diffusion metrics and temporal stability. To achieve this, we compared if the introduction of contrast agent during perfusion, post-fixation or rehydration effects relaxation times or diffusion metrics. By repeatedly scanning the brains over an extended rehydration time we asses both the magnitude and the stability of diffusion and relaxation parameters as a function of the tissue preparation.

By providing a systematic evaluation of contrast agent application effects in the avian brain, this work fills an important methodological gap and extends existing ex-vivo DWI optimization frameworks beyond mammalian models. The protocol presented here is designed to support future high-resolution DWI studies of avian neural circuits and offers a reproducible foundation for comparative connectomics studies across species.

## 2 Methods

### Experimental design and study overview

#### Animals and tissue preparation

For this study a total of 9 pigeons, divided into 4 groups, were perfused to obtain the tissue (Table 1). The animals received an intramuscular injection of 0.1 mL Heparin diluted in 0.1 mL sodium chloride (NaCl) prior to perfusion to prevent blood clots. The animals were then anaesthetized by injection of equithesin (0.45 ml/100 g body weight), after which they were then transcardially perfused with 500 ml of 42 °C warm 0.9 % NaCl, followed by, depending on the experimental group 500 ml of 4 ° C cold 2 % paraformaldehyde (PFA) in 0.12 M phosphate-buffered saline (PBS; pH 7.4) or 500 ml of 4 ° C cold 2 % PFA in 0.12 M PBS containing 15 mM contrast agent. Afterwards the brains were extracted and post-fixed in 2 % PFA for five days.

**Table 1:**
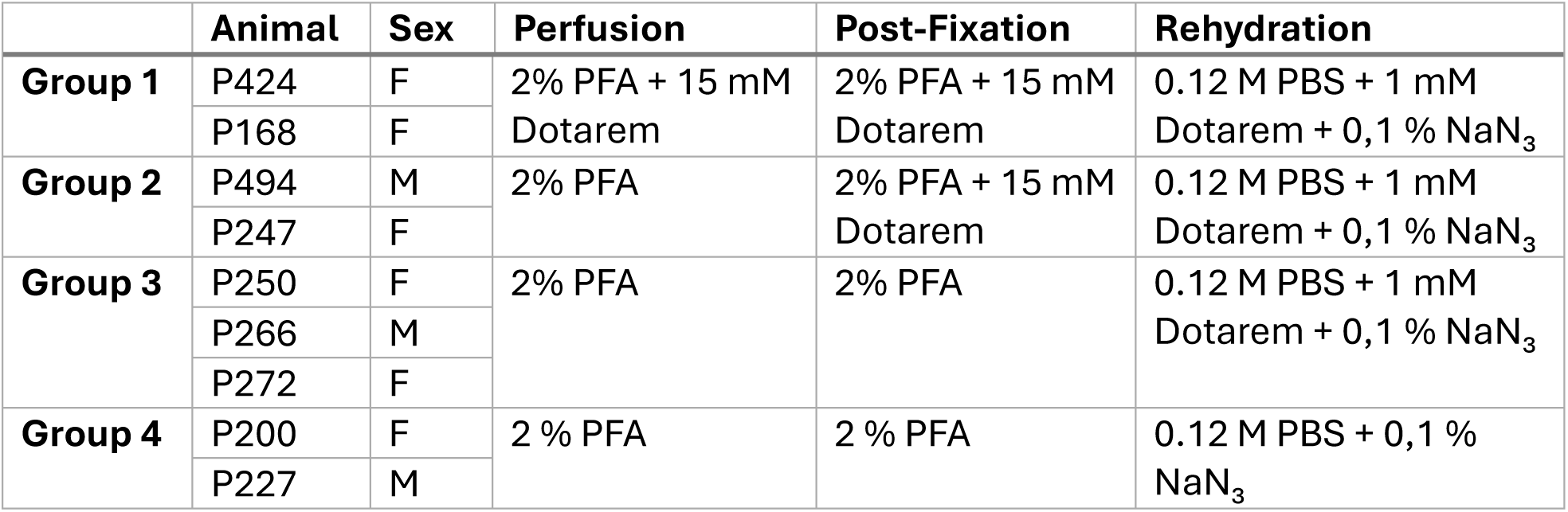
Summary of animals and tissue processing procedures. Animal identification numbers, sex, experimental group assignment, and solutions used during perfusion, post-fixation, and rehydration are listed.

#### Contrast agent application protocols

To investigate the effect of contrast agent on relaxation times and diffusion metrics we applied Dotarem (gadoterate meglumine) as a contrast agent at different stages during the tissue preparation process.

Subjects of the first condition were perfused with 2 % PFA containing 15 mM contrast agent, the brains were then extracted from the skull and subsequently post-fixed in 2 % PFA containing 15 mM contrast agent for 5 days. Subjects of the second condition were perfused with 2 % PFA, after extracting the brains from the skull they were then post-fixed in 2 % PFA containing 15 mM contrast agent for 5 days. Subjects of the third condition were perfused with 2 % PFA and the extracted brains were then post-fixed for 5 days in 2 % PFA. For all three conditions the tissue was rehydrated for 70 days in 1 x PBS containing 1 mM contrast agent and 0,1 % NaN₃ following the post-fixation period. Lastly, subjects of the fourth condition were perfused using 2% PFA, after extracting the brains from the skull the tissue was post-fixed in 2 % PFA for 5 days and subsequently rehydrated in 1 x PBS containing and 0,1 % NaN₃.

For the MRI acquisition of all experimental conditions, brains were positioned in 50 mL centrifuge tubes. To ensure consistent positioning across repeated scans, each brain was fixed to a small plastic support using aquarium-grade adhesive that is insoluble in aqueous solutions. The tubes were filled with the respective rehydration solutions as stated above. Brains were transferred into the tubes immediately prior to the first MRI acquisition, which was designated as Day 0. During the rehydration period, the rehydration solution in all tubes was renewed every 5 days.

#### MRI data acquisition

MRI data was collected using a Bruker BioSpec 7 T scanner (horizontal bore, 70/30 USR, Avance III electronic, Germany) with Paravision 6.0.1 software. RF transmission was done via a quadrature birdcage resonator (82 mm ID), while the quadrature coil was used for data collection.

T1 maps were acquired using a 3D rapid acquisition with relaxation enhancement (RARE) sequence with variable TR = 54, 342, 712, 1231, 2110, 6500 ms, TE = 17.8 ms, RARE factor = 8, one average, acquisition matrix = 60 × 60× 60 and voxel size = 0,43×0,43×0,43 mm. T2 maps were acquired using a multi-slice multi-echo (MSME) sequence with TR = 900 ms, TE = 6, 12, 18, 24, 30, 36, 42, 48, 54, 60, 66, 72, 78, 84, 90, 96, 102, 108, 114, 120, 126, 132, 138, 144 and 150 ms, acquisition matrix = 60×60×60 and voxel size = 0,43×0,43×0,43 mm. T2* maps were acquired using a gradient echo (GRE) sequence with TR = 220 ms, TE = 2, 6, 10, 14, 18, 22, 26, 30, 34, 38, 42, 46, 50, 54, 58, 62, 66, 70, 74, 78, 82, 86, 90, 94, 98, 102, 106, 110, 114 and 118 ms, acquisition matrix = 60×60×60 and voxel size = 0,43×0,43×0,43 mm.

DWI measurement scans were carried out using a 3D diffusion-weighted echo planar imaging (EPI) sequence with a target b-value of 650 s/mm , 40 diffusion directions and 5 non-diffusion-weighted images, TR = 2100 ms, TE = 30,5 ms, four segments, acquisition matrix = 60×60×60 and voxel size = 0,43×0,43×0,43 mm.

### Data Processing

After converting the images from DICOM to NIFTI format using the *dcm2niix* function the images were scaled up by factor 10 to ensure the voxel dimensions match the dimensions from human imaging expected by MRI analysis software.

Firstly, the B0 images were bias field corrected using the *N4BiasFieldCorrection* function, then denoised using the *DenoiseImage* function before extracting the brains using FSL’s *bet* function (Jenkinson et al., 2012; Smith et al., 2004; Woolrich et al., 2009). Lastly the extracted brains were manually corrected. Afterwards, a brain template from the T2 images of each individual was generated using the *antsMultivariateTemplateConstruction.sh* function of the ANTs software (Avants et al., 2011). The T1, T2, and T2* images were skull-stripped using FSL’s *bet* function, before being manually corrected and then registered to the template.

The preprocessing steps for the DWI data were carried out using functionality established in the MRtrix3.0.4 package (Tournier et al., 2019). The DWI data processing steps after converting the images included denoising using Marchenko-Pastur principal component analysis (*dwidenoise*), Gibbs ringing removal (*mrdegibbs*), motion and distortion correction, which included Eddy current distortions and susceptibility-induced EPI distortions (*dwifslpreproc*), and finally bias field correction (*dwibiascorrect*).

To fit the T1, T2 and T2* curves, the built-in relaxation module of the Bruker software was used. To fit T1 the software uses an inversion recovery exponential model applying the following function.

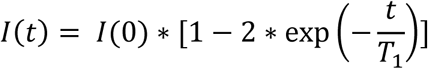

To fit T2 the software uses a mono-exponential decay model applying the following function.

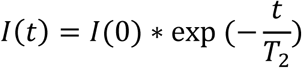

To fit T2* the software also uses a mono-exponential decay model applying the following function.

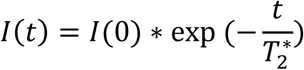

#### Region of Interest

To calculate the mean T1, T2, and T2* values for the nidopallium caudolaterale (NCL), the entopallium (E), and the striatum, a mask for each region of interest (ROI) was manually created in standard template space. The stereotaxic atlas of the pigeon brain by Karten and Hodos (1967) was used as a guide. All ROI masks encompassed the corresponding structures on both hemispheres. The Striatum mask included areas of both the medial striatum (MSt) and the lateral striatum (LSt). The anatomical locations and extents of the ROI masks are illustrated in Figure 1, where the entopallium is shown in red, the striatum in blue, and the NCL in green.

**Figure 1:**
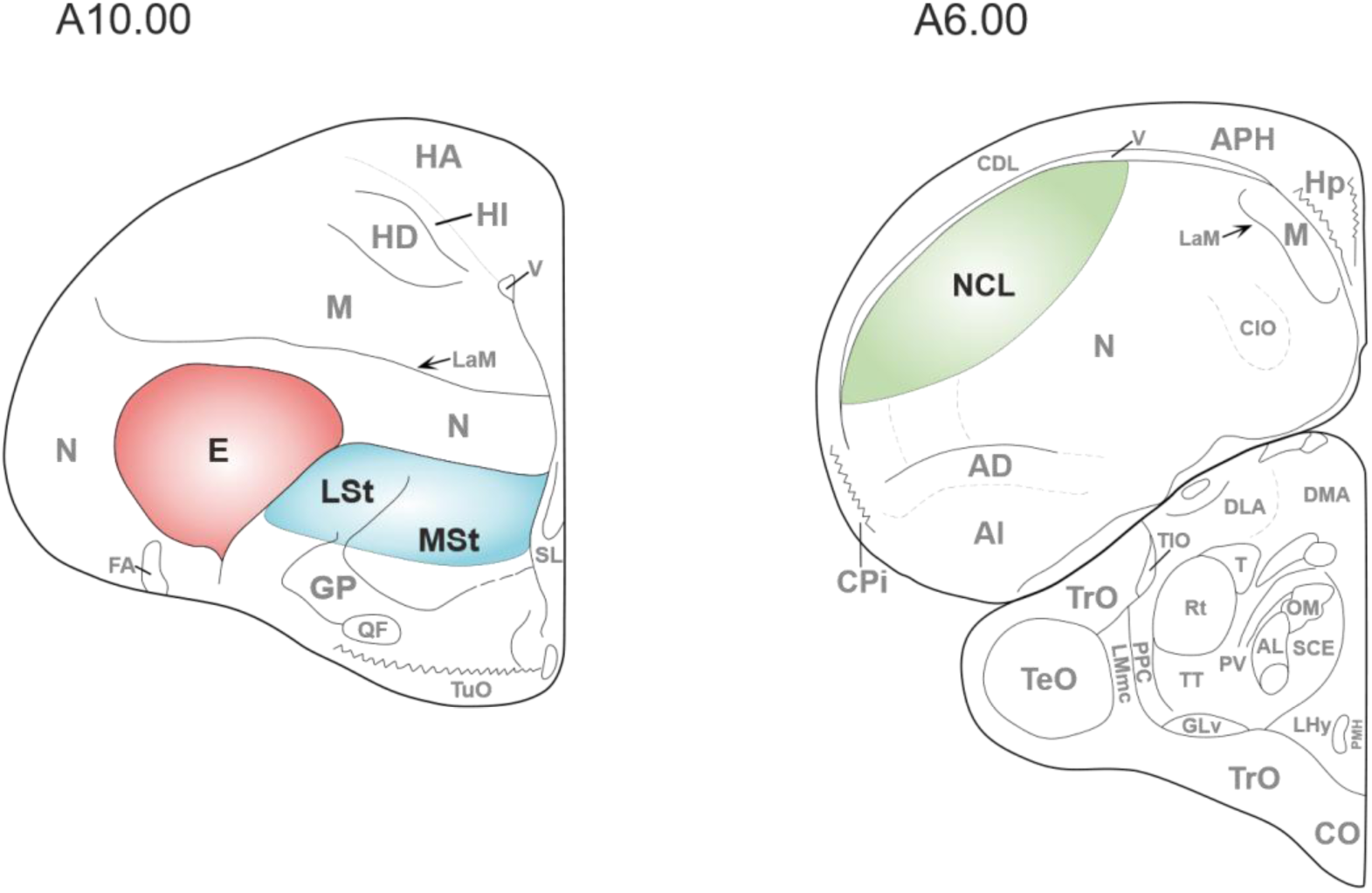
Anatomical localization of regions of interest. Representative atlas sections showing the regions included in the analysis. Colored overlays indicate the entopallium (E, red), Striatum (LSt and MSt, blue), and nidopallium caudolaterale (NCL, green). Boundaries are shown schematically and represent the areas sampled for quantification.

The masks were then registered to the T1, T2 and T2* images. Custom MATLAB scripts (v2025a) were used to calculate the means and standard errors for all three images, all ROIs and all individuals.

#### Diffusion tensor fitting

The pre-processed and corrected data was then diffusion fitted using FSL’s DTIFIT. The FA maps and MD maps were part of the functions output and subsequently the maps were registered to the template

The strategy to calculate the mean FA and MD values in the NCL, the entopallium and the striatum was identical to how the means of the relaxation times were determined.

## 3 Results

### 3.1 Effects of contrast-agent application protocol on relaxation times

To assess if adding contrast agent to the fixative during the perfusion and post-fixation we measured the relaxation times at different time points during the rehydration phase. The scans were performed at days 0, 6, 13, 22 and 70 of the rehydration period. We then calculated the mean values and the standard error of the mean for T1, T2 and T2* in the Entopallium, Striatum, and NCL. The values of all parameters, for each animals and scan day are displayed for the Entopallium in Table 2, for the Striatum in Table 3 and for the NCL in Table 4.

**Table 2:**
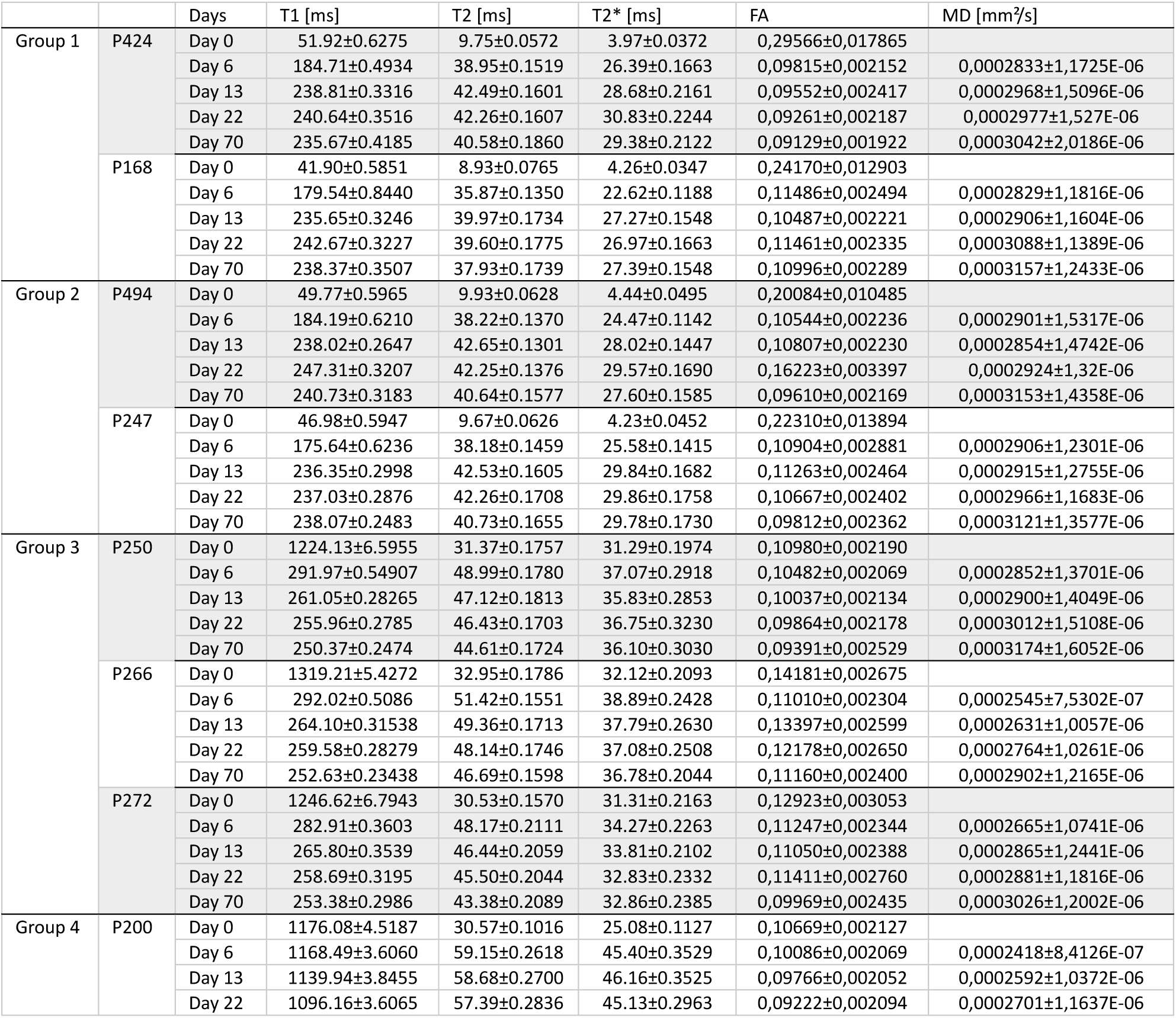
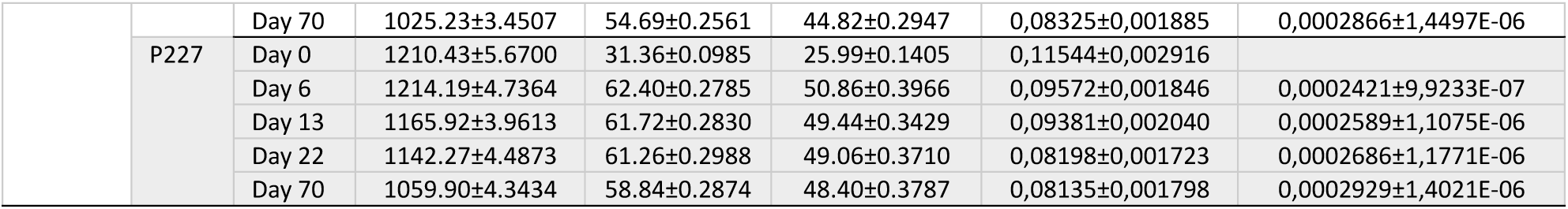
Quantitative MRI parameters of the Entopallium across experimental groups and time points. Mean ± standard error values of T1, T2, T2*, fractional anisotropy (FA), and mean diffusivity (MD) measured in the NCL for each animal and time point

**Table 3:**
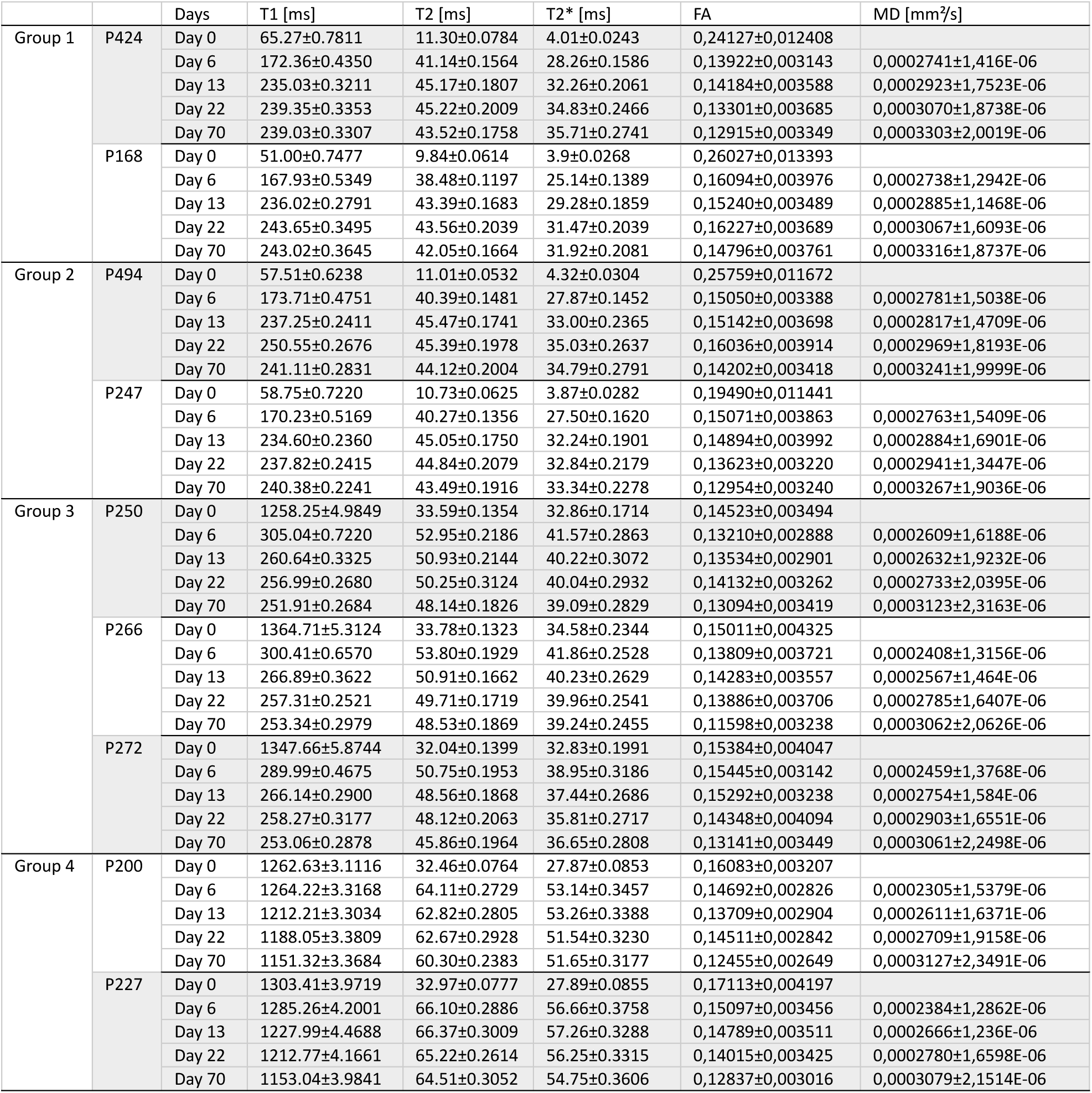
Quantitative MRI parameters of the Striatum across experimental groups and time points. Mean ± standard error values of T1, T2, T2*, fractional anisotropy (FA), and mean diffusivity (MD) measured in the NCL for each animal and time point

**Table 4:** Quantitative MRI parameters of the nidopallium caudolaterale (NCL) across experimental groups and time points. Mean ± standard error values of T1, T2, T2*, fractional anisotropy (FA), and mean diffusivity (MD) measured in the NCL for each animal and time point

|  |  | Days | T1 [ms] | T2 [ms] | T2* [ms] | FA | MD [mm<sup>2</sup>/s] |
| --- | --- | --- | --- | --- | --- | --- | --- |
| Group 1 | P424 | Day 0 | 49.00±0.9469 | 10.38±0.0870 | 4.84±0.0295 | 0,29342±0,016729 | |
| | | Day 6 | 204.23±0.4289 | 45.55±0.4429 | 29.16±0.6934 | 0,08114±0,001720 | 0,0002533±2,2927E-06 |
| | | Day 13 | 237.50±0.2689 | 47.39±0.2086 | 31.59±0.2044 | 0,07059±0,001743 | 0,0002611±3,1207E-06 |
| | | Day 22 | 236.61±0.3160 | 48.78±0.4049 | 33.77±0.3434 | 0,07335±0,001912 | 0,0002794±2,9321E-06 |
| | | Day 70 | 232.51±0.3131 | 45.59±0.1746 | 34.48±0.2433 | 0,06880±0,001804 | 0,0002980±2,9017E-06 |
| | P168 | Day 0 | 36.44±0.4662 | 10.09±0.0738 | 5.50±0.0395 | 0,27966±0,013921 | |
| | | Day 6 | 193.27±0.9423 | 40.56±0.2179 | 23.89±0.1848 | 0,10249±0,002613 | 0,0002427±2,4152E-06 |
| | | Day 13 | 231.96±0.2575 | 45.96±0.1935 | 29.33±0.1874 | 0,07594±0,001775 | 0,0002505±2,9815E-06 |
| | | Day 22 | 238.35±0.2265 | 46.96±0.3274 | 31.76±0.2339 | 0,08958±0,001712 | 0,0002733±2,4823E-06 |
| | | Day 70 | 236.33±0.2487 | 45.35±0.1966 | 32.23±0.2077 | 0,07363±0,001619 | 0,0002994±2,5987E-06 |
| Group 2 | P494 | Day 0 | 41.70±0.4612 | 10.73±0.0818 | 5.96±0.0377 | 0,18621±0,011779 | |
| | | Day 6 | 208.31±0.5016 | 44.47±0.2655 | 27.38±0.1627 | 0,08117±0,001727 | 0,0002635±2,9641E-06 |
| | | Day 13 | 232.41±0.1766 | 46.81±0.2012 | 31.92±0.1506 | 0,07906±0,001636 | 0,0002512±3,2581E-06 |
| | | Day 22 | 238.27±0.1991 | 45.80±0.1801 | 33.09±0.1830 | 0,12615±0,002057 | 0,0002765±2,7272E-06 |
| | | Day 70 | 237.07±0.2245 | 44.39±0.1727 | 32.68±0.1720 | 0,07940±0,001629 | 0,0002956±3,1542E-06 |
| | P247 | Day 0 | 41.34±0.8308 | 10.32±0.0788 | 5.05±0.0299 | 0,27135±0,016187 | |
| | | Day 6 | 190.07±0.5416 | 42.48±0.2510 | 26.08±0.2267 | 0,07666±0,001881 | 0,0002442±2,8795E-06 |
| | | Day 13 | 230.86±0.1965 | 46.31±0.2365 | 30.89±0.2072 | 0,07429±0,001751 | 0,0002610±2,8149E-06 |
| | | Day 22 | 230.87±0.1936 | 46.31±0.2966 | 31.64±0.2343 | 0,08978±0,001957 | 0,0002365±3,4775E-06 |
| | | Day 70 | 233.49±0.1712 | 45.34±0.2981 | 31.79±0.2385 | 0,07122±0,001704 | 0,0002602±4,1252E-06 |
| Group 3 | P250 | Day 0 | 1388.36±6.0253 | 39.65±0.2929 | 41.63±0.4101 | 0,13619±0,004934 | |
| | | Day 6 | 281.58±0.4161 | 59.13±0.2968 | 44.92±0.3135 | 0,06556±0,001372 | 0,0002179±3,4879E-06 |
| | | Day 13 | 261.44±0.3493 | 55.79±0.2500 | 43.33±0.3066 | 0,06242±0,001375 | 0,0002316±3,0503E-06 |
| | | Day 22 | 256.64±0.3018 | 58.99±0.7011 | 44.36±0.4389 | 0,06859±0,001523 | 0,0002381±2,9214E-06 |
| | | Day 70 | 254.00±0.2693 | 53.33±0.2770 | 41.70±0.2761 | 0,06127±0,001405 | 0,0002643±2,7125E-06 |
| | P266 | Day 0 | 1467.97±6.1402 | 40.45±0.3057 | 38.59±0.3232 | 0,10359±0,003169 | |
| | | Day 6 | 279.36±0.4383 | 58.88±0.2919 | 42.94±0.2978 | 0,07968±0,001918 | 0,0002207±1,5568E-06 |
| | | Day 13 | 263.61±0.2598 | 56.17±0.2090 | 42.50±0.2914 | 0,07434±0,001601 | 0,0002349±2,0754E-06 |
| | | Day 22 | 257.60±0.2601 | 54.23±0.1622 | 42.76±0.2811 | 0,07551±0,001682 | 0,0002473±1,8492E-06 |
| | | Day 70 | 256.37±0.2342 | 53.00±0.2364 | 41.74±0.3084 | 0,07140±0,001636 | 0,0002695±2,4739E-06 |
| | P272 | Day 0 | 1431.53±6.5665 | 38.18±0.3125 | 37.79±0.4159 | 0,19274±0,012758 | |
| | | Day 6 | 275.98±0.4199 | 56.59±0.2267 | 43.04±0.2384 | 0,07721±0,001570 | 0,0002094±3,046E-06 |
| | | Day 13 | 265.49±0.3280 | 54.31±0.2404 | 41.85±0.2787 | 0,08024±0,001728 | 0,0002377±2,5412E-06 |
| | | Day 22 | 257.11±0.2473 | 53.47±0.2574 | 40.76±0.2711 | 0,08402±0,001925 | 0,0002464±2,4751E-06 |
| | | Day 70 | 255.69±0.2728 | 50.45±0.2099 | 40.32±0.2444 | 0,06733±0,001508 | 0,0002682±2,946E-06 |
| Group 4 | P200 | Day 0 | 1487.93±3.4176 | 38.94±0.2907 | 32.78±0.2369 | 0,09399±0,001916 | |
| | | Day 6 | 1414.42±2.6821 | 73.23±0.2464 | 65.52±0.4332 | 0,07806±0,001712 | 0,0002515±1,0031E-06 |
| | | Day 13 | 1374.08±2.7238 | 71.21±0.2491 | 63.92±0.3195 | 0,08946±0,001937 | 0,0002669±1,2048E-06 |
| | | Day 22 | 1339.48±2.9368 | 69.51±0.2302 | 61.58±0.2405 | 0,07655±0,001542 | 0,0002816±1,2513E-06 |
| | | Day 70 | 1291.63±3.0913 | 67.76±0.2201 | 60.56±0.2467 | 0,07099±0,001492 | 0,0003064±1,5292E-06 |
| | P227 | Day 0 | 1538.34±6.3676 | 43.84±0.5097 | 35.73±0.3128 | 0,09596±0,002513 | |
| | | Day 6 | 1469.08±4.4568 | 79.65±0.4711 | 69.88±0.5942 | 0,07215±0,001597 | 0,0002230±1,3568E-06 |
| | | Day 13 | 1435.89±7.8287 | 77.82±0.5642 | 65.93±0.4330 | 0,07470±0,001631 | 0,0002366±1,4581E-06 |
| | | Day 22 | 1377.61±5.2029 | 73.82±0.3769 | 65.01±0.4065 | 0,07459±0,001461 | 0,0002480±1,8245E-06 |
| | | Day 70 | 1342.57±6.4335 | 75.33±0.5629 | 68.36±0.6259 | 0,05502±0,001187 | 0,0002854±1,5846E-06 |

T1 relaxation times showed protocol-dependent longitudinal changes with the largest effects being observed with regards to if contrast agent was added during tissue fixation and rehydration or exclusively during tissue rehydration (Figure 2).

**Figure 2:**
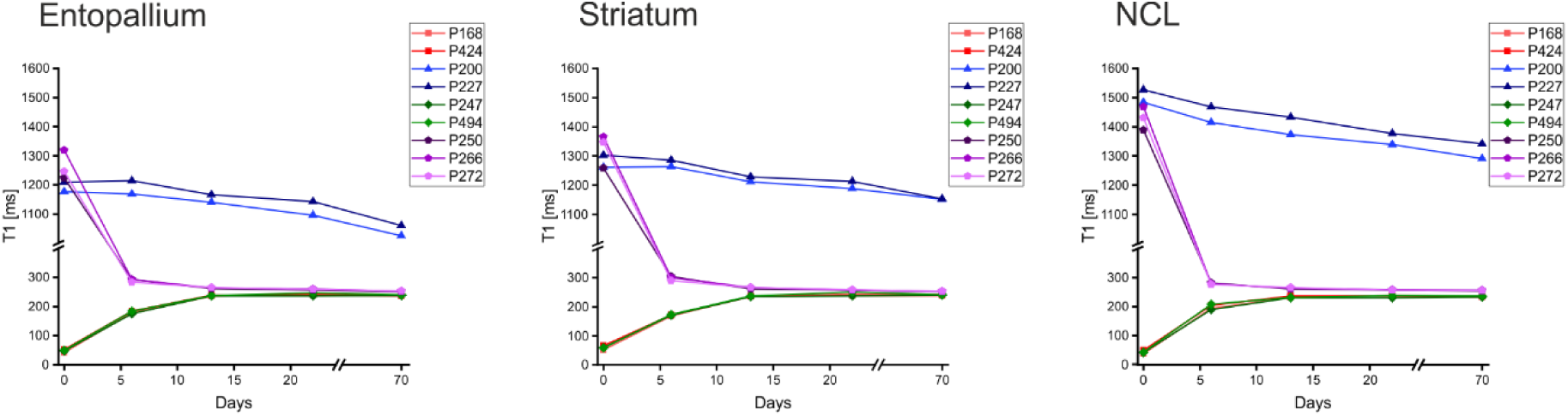
Mean T1 value ± standard error of the mean in the Entopallium, Striatum and NCL. Animals of group 1 (P424, P168) are depicted in red, animals of group 2 (P494, P247) in green, animals of group 3 (P250, P266, P272) in purple and animals of group 4 (P200, P227) in blue.

In the groups that had contrast agent added to the fixative during any step of tissue fixation an increase in T1 can be observed from day 0 to day 13. At day 0 the mean T1 values range from 41.90 ms – 51.92 ms in the Entopallium, from 51.00 ms – 65.27 ms in the Striatum and from 36.44 ms – 49.00 ms in the NCL. By day 13 they have increased to 235.65 ms – 238.81 ms in the Entopallium, to 234.60 ms – 237.25 ms in the Striatum and to 230.86 ms – 237.50 ms in the NCL. Afterwards the values seem to be rather stable in all areas and individuals of these conditions. From day 13 to day 70 the T1 values change 0.7 % - 1.3 % in the Entopallium, 1.6 % - 2.9 % in the Striatum and1.1 % - 2.1 % in the NCL.

When contrast agent was only added during the rehydration phase, T1 values decreased substantially from Day 0 to Day 13. In the entopallium, mean T1 decreased from 1388.36 ms – 1467.97 ms at Day 0 to 261.44 ms – 265.49 ms at Day 13. Similarly, mean T1 values in the Striatum decreased from 1258.25 ms – 1364.71 ms to 260.64 ms – 266.89 ms while mean T1 values in the NCL decreased from 1388.36ms – 1467.97 ms to 261.44 ms – 265.49 ms. Similar to the prior conditions described, the T1 values remain relatively stable from day 13 onwards until day 70. During this time a decrease of 4 % – 4.6 % can be reported in the Entopallium, a decrease of 3.3 % - 5.1 % in the Striatum and a decrease of 2.7 % - 3.7 % in the NCL.

In contrast, the tissue with no exposure to contrast agent at any time seem to show a slight but steady decrease of T1 values of 8.8 % - 13.2 % across all observed areas over the 70-day rehydration period.

T2 relaxation times also show protocol-dependent changes, most notable is that T2 is highest when no contrast agent was added during any tissue preparation step (Figure 3).

**Figure 3:**
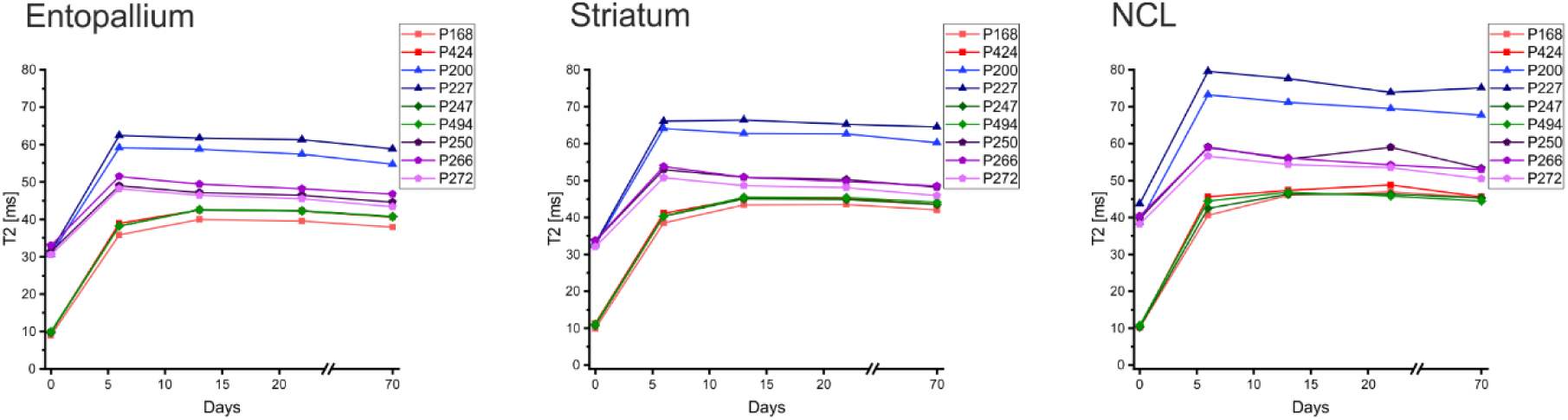
Mean T2 values ± standard error of the mean in the Entopallium, Striatum and NCL. Animals of group 1 (P424, P168) are depicted in red, animals of group 2 (P494, P247) in green, animals of group 3 (P250, P266, P272) in purple and animals of group 4 (P200, P227) in blue.

When contrast agent was added during either the perfusion, the post-fixation and the rehydration phase or only during the post-fixation and the rehydration phase, T2 values at day 0 of the rehydration phase are relatively low in all areas. The mean values are 8.93 ms – 9.93 ms in the Entopallium, 9.84 ms – 11.30 ms in the Striatum and 10.09 ms – 10.73 ms in the NCL, at day 0. From day 0 to day 13 they increase to 39.97 ms – 42.65 ms in the Entopallium, 43.39 ms – 45.47 ms in the Striatum and to 45.96 ms – 47.39 ms in the NCL. Subsequently the values then decrease slightly from day 13 to day 70. Decreases of 4.2 % - 5.1 % can be observed in the Entopallium, 3.0 % - 3.6 % in the Striatum and 1.3 % - 5.2 % in the NCL.

In contrast, when the tissue was not exposed to contrast agent prior to day 0 of the rehydration phase higher T2 values can be reported at this point in time for these groups. The mean T2 values for the Entopallium were 30.53 ms – 32.95 ms, for the Striatum 32.04 ms – 33.78 ms and for the NCL 38.18 ms – 43.84 ms. When contrast agent was introduced to the tissue during the rehydration phase, we then see an increase in T2 values from day 0 to day 6 which is followed by a slight decrease from day 6 to day 70. In the Entopallium the values increase to 48.17 ms – 51.42 ms from day 0 to day 6 and then decrease to 43.38 ms - 46.69 ms from day 6 to day 70. In the Striatum the mean T2 values increase from day 0 to day 6 to 50.75 - 53.80 ms, from day 6 to day 70 they then decrease to 45.86 – 48.53 ms. In the NCL the values increase to 58.88 – 59.13 ms from day 0 to day 6 followed by a decrease to 50.45 – 53.33 ms from day 6 to day 70.

For the control group, that had no contrast agent introduced at any point during tissue preparation, the T2 values also increased from day 0 to day 6 and then decreased from day 6 to day 70. The values increase to 59.15 ms – 62.40 ms from day 0 to day 6 and then decrease to 54.69 ms – 58.84 ms from day 6 to day 70 in the Entopallium. In the Striatum the values increase from day 0 to day 6 to 64,11 ms - 66,10 ms and then decrease from day 6 to day 70 to 60.30 ms – 64.51 ms. In the NCL we can observe an increase to 73.23 ms – 79.65 ms from day 0 to day 6 followed by a decrease to 67.76 ms – 75.33 ms from day 6 to day 70.

The changes in T2* show a pattern similar to the T2 values, with an initial increase in all groups; however, T2* values then either change only slightly or remain stable over time (Figure 4).

**Figure 4:**
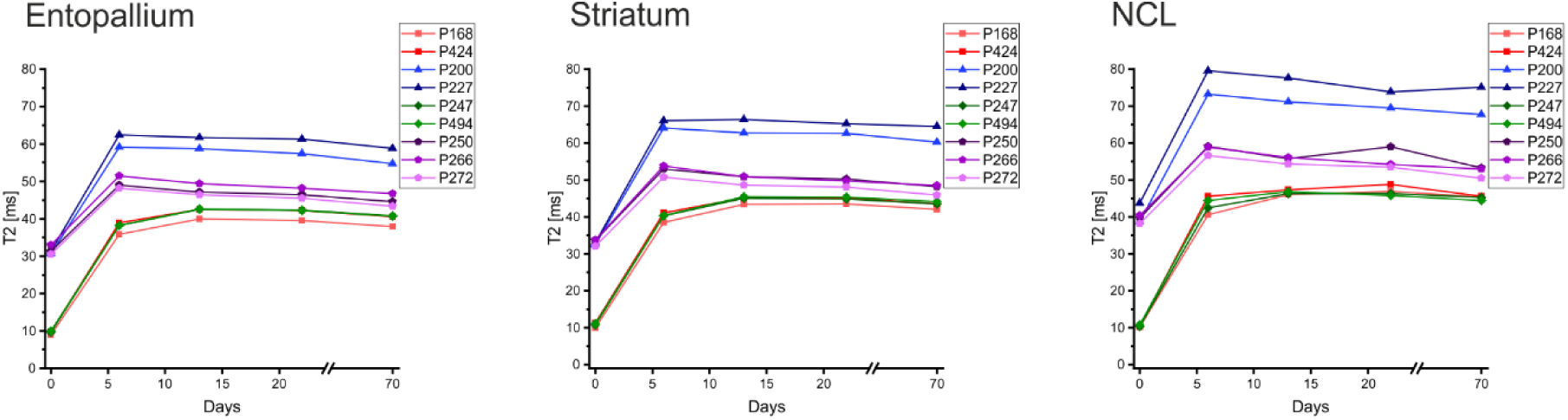
Mean T2* values ± standard error of the mean in the Entopallium, Striatum and NCL. Animals of group 1 (P424, P168) are depicted in red, animals of group 2 (P494, P247) in green, animals of group 3 (P250, P266, P272) in purple and animals of group 4 (P200, P227) in blue.

**Figure 5:**
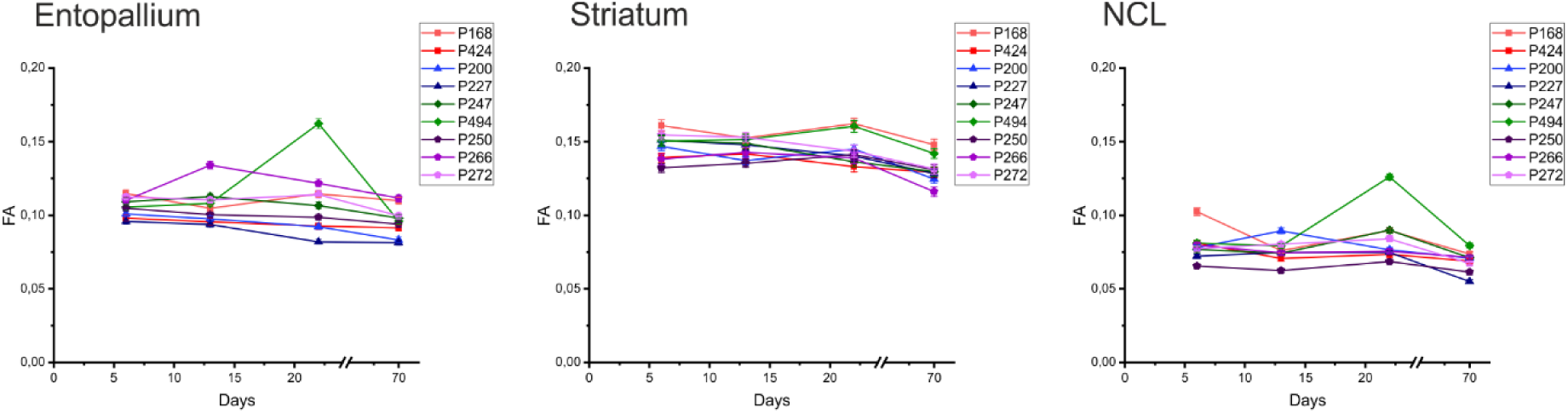
Average Fractional Anisotropy (FA) values +-standard error of the mean in the Entopallium, Striatum and NCL. Animals of group 1 (P424, P168) are depicted in red, animals of group 2 (P494, P247) in green, animals of group 3 (P250, P266, P272) in purple and animals of group 4 (P200, P227) in blue.

**Figure 6:**
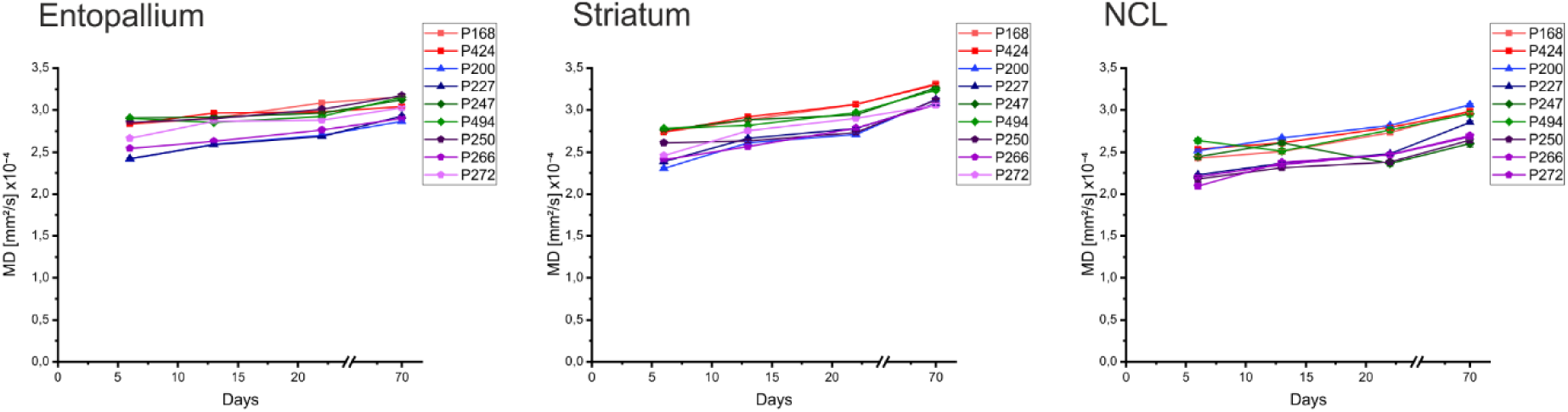
Average Mean Diffusivity (MD) values +-standard deviation in the Entopallium, Striatum and NCL. Animals of group 1 (P424, P168) are depicted in red, animals of group 2 (P494, P247) in green, animals of group 3 (P250, P266, P272) in purple and animals of group 4 (P200, P227) in blue.

When contrast agent was added during either the perfusion, the post-fixation and the rehydration phase or only during the post-fixation and the rehydration phase, T2* values at day 0 of the rehydration phase are relatively low across all areas in the respective groups. The mean T2* values are 3.97 ms – 4.44 ms for the Entopallium, 3.87 ms – 4.32 ms for the Striatum and 4.84 ms – 5.96 ms for the NCL at day 0. From day 0 to day 6, an increase to 22.62 ms to 26.39 ms in the Entopallium, 25.14 ms – 28.26 ms in the Striatum and 23.89 ms – 29.16 ms in the NCL can be observed. Subsequently, the values then change slightly from day 6 on and at day 70 the mean T2* values are 27.39 ms – 29.78 ms for the Entopallium, 31.92 ms – 35.71 ms for the Striatum and 31.79 ms – 34.48 ms for the NCL.

In contrast, when the tissue was not exposed to contrast agent prior to day 0 of the rehydration phase, higher T2* values are observed in these groups at this time point. The mean T2* values for the Entopallium are 25.08 ms – 32.12 ms, for the Striatum 27.87 ms – 34.58 ms and for the NCL 32.78 ms – 41.63 ms. When contrast agent is introduced during the rehydration phase, an increase in T2* values from day 0 to day 6 is observed, followed by a slight decrease from day 6 to day 70. In the Entopallium, mean T2* values increase to 34.27 ms - 38.89 ms from day 0 to day 6 and then decrease to 32.86 ms – 36.78 ms from day 6 to day 70. In the Striatum, values increase from day 0 to day 6 to 38.95 ms – 41.86 ms, and then decrease from day 6 to day 70 to 36.65 ms – 39.24 ms In the NCL, values increase to 42.94 ms – 44.92 ms from day 0 to day 6, followed by a decrease to 40.32 ms – 41.74 ms from day 6 to day 70.

For the control group, in which no contrast agent was introduced at any point during tissue preparation, T2* values also increase from day 0 to day 6 and then decrease from day 6 to day 70. In the Entopallium, values increase to 45.40 ms – 50.86 ms from day 0 to day 6 and then decrease to 44.82 ms – 48.40 ms from day 6 to day 70. In the Striatum, values increase from day 0 to day 6 to 53.14 ms – 27.89 ms, and then decrease from day 6 to day 70 to 51.65 ms – 54.75 ms. In the NCL, an increase to 65.52 ms – 69.88 ms from day 0 to day 6 is observed, followed by a decrease to 60.56 ms – 68.36 ms from day 6 to day 70.

### 3.2 Impact of contrast-agent protocol on diffusion tensor metrics

Furthermore, to assess the impact of the contrast agent on diffusion metrics we calculate the average fractional anisotropy and average mean diffusivity in the entopallium, the striatum and the NCL. For these measures we only analyzed the scan days 6,13,22 and 70, as on day 0 T2 relaxation times were too low to extract accurate FA and MD values. The values of all parameters, for each animals and scan day are displayed for the Entopallium in Table 2, for the Striatum in Table 3 and for the NCL in Table 4.

While the FA values differ across the regions of interest, within each region the changes over time are minimal. No protocol-dependent changes can be clearly observed. In general, we find that for the regions we analyzed FA is highest in the Striatum and lowest in the NCL.

Furthermore, we also do not observe protocol-dependent changes in the average mean diffusivity values across all areas. In general, we consistently across all animals and areas see a slight increase of the MD values from day 6 to day 70.

## 4 Discussion

### 4.1 Contrast agent timing as a key determinant of relaxation behavior

This study shows that the timing of contrast agent exposure is a primary determinant of longitudinal relaxation behavior in ex vivo avian neural tissue. Across all brains areas we analyzed, a distinct temporal change can be observed, that depended on whether contrast agent was introduced during the tissue fixation or exclusively during the rehydration phase.

When tissue was exposed to contrast agent during the perfusion and/or post-fixation, T1 values showed a pronounced increase during the first two weeks of the rehydration period. It was shown that gadolinium-based contrast agents significantly lower T1 relaxation times (Rogosnitzky & Branch, 2016). Thus, the observed increase in T1 seems to be driven by the change in contrast agent concentration in the tissue. Although the tissue was exposed to 15 mM contrast agent during fixation, the concentration was reduced to only 1 mM during the rehydration phase; consequently, the negative effects of the contrast agent were less pronounced during rehydration. After this initial increase the T1 relaxation time stabilized and stayed consistent throughout the rest of the rehydration period.

In contrast, when contrast agent was introduced only during the rehydration phase, T1 values show a strong initial decrease, driven by the effect of contrast agent on relaxation times, before converging to similar stable levels by day 13 to 22. To summarize, all contrast agent treated conditions ultimately reached comparable equilibrium T1 values, at a similar time. This indicates that the final longitudinal relaxation behavior is largely independent of the initial application timing of the contrast agent. This convergence strongly suggests that diffusion-driven equilibration of the contrast agent within the tissue is the dominant mechanism governing long-term T1 relaxation behavior.

A similar pattern was observed for T2 and T2*, where all conditions with contrast agent exposure during tissue fixation showed initially low values which then increased and finally stabilized. This initial increase is probably mainly driven by washing unbound fixative out of the tissue (Shepherd et al., 2009) and to a smaller extend by the reduction in contrast agent concentration (Rogosnitzky & Branch, 2016). When the tissue was not exposed to contrast agent prior to the rehydration phase we observed higher values at the initial scan, yet we still observe an increase in T2 and T2* from day 0 to day 6. However, the values for these two groups then diverged during the rehydration period. The tissue that had contrast agent introduced during this stage exhibited lower T2 and T2*, indicating the negative effect contrast agent has on T2 and T2* relaxation time. As the initial increase in transverse relaxation time is present in all groups, including the control group, it seems to stem from removing unbound PFA from the tissue (Shepherd et al., 2009; Ullmann et al., 2010).

Contrary to what we observed for the T1 relaxation time, the contrast agent exposure during tissue fixation does seem to have a slight negative, long-lasting and irreversible effect of T2 and T2*relaxation behavior of the tissue.

In general, it should be noted that a recent study investigating tissue optimization for high-quality ex vivo diffusion MRI in rats (Barrett et al., 2023) found that the inclusion of contrast agent in the perfusate resulted in lower T1 values even after the rehydration period.

Although all samples were subsequently rehydrated in a 1 mM contrast agent solution, tissues exposed to contrast agent during perfusion retained lower T1 values than tissues perfused without contrast agent. While we cannot comment on potential differences in T1 values arising from the contrast agent application protocol, it should be noted that the study used Magnevist (Gd-DTPA), whereas we used Dotarem (Gd-DOTA). Therefore, it remains unclear to what extent the observed differences between studies reflect the perfusion protocol, the contrast agent itself, or an interaction between both factors. Furthermore, species-specific differences in tissue composition, fixation response, or contrast agent distribution cannot be excluded, given that the previous study was conducted in rats while our study investigated pigeon brains.

### 4.2 Preservation of diffusion metrics across preparation protocols

Despite the differences of contrast agent application on tissue relaxation, diffusion tensor metrics remained stable across all conditions. Neither the fractional anisotropy nor the mean diffusivity showed a systematic difference between contrast agent protocols, and temporal changes within each protocol were minimal. These findings are rather consistent with recent studies on the topic in mammalian brains (Barrett et al., 2023)

This difference between relaxation behavior and diffusion metrics is particularly important, as it indicates that contrast agent application, when used at the tested concentration, does not substantially alter microstructural measurements. This stability of fractional anisotropy across time and conditions suggests that the fiber organization is preserved regardless of contrast agent exposure.

Regarding MD, we observed a slight increase over time across all conditions, suggesting a global tissue effect rather than a contrast-agent-specific mechanism. One possible explanation is ongoing tissue rehydration and equilibration following fixation, which may partially alleviate fixation-induced diffusion restrictions (Roebroeck et al., 2019; Shepherd et al., 2009).

Importantly, the absence of protocol-dependent differences in diffusion metrics suggests that the primary role of contrast agent in this context is to modulate relaxation properties and improve SNR efficiency, rather than altering the diffusion signal itself. This supports the use of contrast agent enhanced ex vivo imaging protocols for high-resolution DWI without compromising quantative diffusion analysis.

### 4.3 Implications for ex-vivo diffusion imaging in avian models

The results of this study can have direct methodological implications for ex vivo DWI in avian brains and, more broadly, for comparative neuroimaging.

Most notable, we show that contrast agent application solely during the rehydration phase is sufficient to achieve stable relaxation parameters. From a practical perspective, restricting contrast agent application to the rehydration phase offers several advantages including a reduced protocol complexity and improved experimental flexibility. And potentially most importantly the reduced risk of contrast agent overexposure, which could otherwise lead to excessively shortened T2 and T2* (Barrett et al., 2023; Ullmann et al., 2010).

Moreover, the observed convergence of relaxation parameters across protocols suggests that standardization efforts in ex vivo DWI should prioritize sufficient equilibration times of 2 to 3 weeks rather than contrast agent application during tissue fixation. Importantly these findings extend existing optimization frameworks, which are largely derived from mammalian models to the avian brain. Given the substantial anatomical and microstructural differences between birds and mammals, this study provides a necessary species-specific validation of tissue preparations for ex-vivo DWI.

### 4.4 Limitations and future directions

It should be noted that this study has several limitations. Only a single contrast agent concentration was tested. Although the selected concentration was based on findings from a previous study (Barrett et al., 2023) and yielded robust and stable results, a systematic investigation of different contrast agent concentrations could further optimize the balance between T1 shortening and the preservation of T2 and T2* values, thereby helping to identify optimal concentration ranges for different imaging applications. Furthermore, it should be noted that this study tested only a single contrast agent. Therefore, we cannot exclude the possibility that other contrast agents may yield more favorable relaxation metrics. Also, while we assessed major pallial and subpallial regions, finer anatomical structures and specific fibre tracts were not analyzed separately in this study. Given the heterogenous organization of the avian brain, future work could investigate whether smaller or more specialized structures show differential sensitivity to preparation protocols. Finally, this study focused exclusively on ex-vivo conditions. Although the findings are highly relevant for high-resolution structural imaging, the relationship between ex-vivo diffusion properties and in-vivo microstructure remain an important open question, particularly in comparative contexts.

### 4.5. Conclusion

In summary, this study demonstrates that contrast agent timing primarily affects early relaxation dynamics while it only has a minimal impact on long-term equilibrium states and diffusion tensor metrics in ex-vivo avian brain tissue.

We show that T1 relaxation times converge across protocols after approximately 2 to 3 weeks regardless of if contrast agent was added during tissue fixation or solely during the rehydration phase. We also show that adding contrast agent during tissue fixation seems to have a long-lasting slight negative effect on T2 and T2* relaxation times. Furthermore, our findings suggest that diffusion metrics remain robust across all tissue preparation strategies.

Together, these findings establish a simplified, reproducible, and effective tissue preparation protocol for ex vivo DWI in the avian brain. This work provides a methodological foundation for future high-resolution studies of avian brain connectivity and supports the broader goal of standardized comparative connectomics across species.

## Acknowledgement

This work was supported by the Deutsche Forschungsgemeinschaft (DFG; Project No. 395940726 / SFB1372 to OG) and by the European Research Council (ERC) Advanced Grant “AVIAN MIND - Inquiries into a Different Kind of Mind”, project no. 101021354.

## Notes

### Competing Interest Statement

The authors have declared no competing interest.

